# An induced pluripotent stem cell-based chemical genetic approach for studying spinal muscular atrophy

**DOI:** 10.1101/2025.11.04.686319

**Authors:** Richard M. Giadone, Kristina M. Holton, Xiaoyu Hu, Ted Natoli, Sabrina Ghosh, Stanley P. Gill, Aravind Subramanian, Lee L. Rubin

## Abstract

Spinal muscular atrophy (SMA) is a genetic disease characterized by degeneration of spinal cord motor neurons and neuromuscular junctions. Despite recent development in therapies for SMA, treatment efficacy largely relies on administration of drugs early in disease progression and is impacted by underlying patient genetics. Drug discovery for other diseases of the central nervous system (CNS) has also been hindered by heterogeneity in patient genetics and clinical presentations, as well as the need for early intervention. To address these hurdles, we utilized a chemical genetic-based screening approach to adapt the Connectivity Map (CMAP)/L1000 platform to study SMA. To do this, we differentiated moderate and severe SMA patient-specific induced pluripotent stem cells into neuronal cells utilizing a forward programming differentiation protocol, exposed each to 360 neuroactive or CNS disease-related compounds, and interrogated resulting changes in expression of >400 neural genes in a platform we term CMAP_neuro_. In doing so, we generated 4,559 transcriptional profiles identifying stimuli that modulate gene expression differences across SMA neurons. Finally, we make these data queryable, allowing the research community to 1.) identify CNS disease-related perturbagens that mimic or reverse differentially expressed genes, or 2.) explore the transcriptional response of a given perturbation in diverse SMA neuronal cells.

## Introduction

Spinal muscular atrophy (SMA) is a degenerative motor neuron disease caused by a deficiency of Survival of Motor Neuron (SMN) protein resulting from mutations in the *SMN1* gene^1,2^. In patients, disease severity is mainly, though not exclusively, determined by copy number of the truncated, less functional *SMN2* gene, with clinical features classified by the age of detectable changes in motor symptoms and maximum motor function achieved^1^. Two currently approved therapies for SMA modulate splicing of the *SMN2* transcript to increase SMN levels, while a third acts via viral delivery of the *SMN1* gene to patient cells^2–4^. For each, efficacy relies on administration of drug early in disease progression – a problem exacerbated when considering frequently late diagnoses, particularly in milder forms of the disease^5^. To complement these strategies, new approaches seek to identify additional drugs capable of rescuing phenotypic changes resulting from low SMN levels in conjunction with therapeutics to raise SMN levels^6^.

Historically, drug discovery approaches for central nervous system (CNS) diseases have proven challenging in part due to heterogeneity of genetics and patient presentation. Adding to this complexity, a majority of these disorders have multiple traits and phenotypes with unclear linkage between genetic risk loci and functional consequences^7–9^. At the same time, cellular models for CNS disorders are limited by the inability of cultured cells to replicate disease physiology. To address these limitations, several studies have utilized potent stressors such as the proteasome inhibitor MG132 or ER stressors such as thapsigargin or tunicamycin to exacerbate *in vitro* phenotypes for CNS disorders^10,11^. Although successful in this regard, such robust stimuli may mask individual, physiologically relevant cellular and genetic variation, thereby masking more subtle differences caused by disease-associated genes. Building on these approaches, we posit that studying the response of patient-specific cells to large numbers of diverse perturbagens may uncover novel aspects of CNS disease biology.

Fortuitously, novel chemical genetic approaches for drug discovery have coincided with technological advances in high-throughput screening and gene expression profiling platforms^12,13^. In one notable example, the Connectivity Map (CMAP) allows for the discovery of functional connections between diseases, genes, and drugs by combining comprehensive perturbation databases with transcriptomic profiles of cancer cell lines treated with thousands of perturbagens^12,13^. Further, the Luminex1000 (L1000), a cost-effective, high-throughput bead-based platform, is employed within the CMAP pipeline to measure the expression profile of approximately 1000 landmark genes and impute expression of the remaining genes in the transcriptome. To date, CMAP includes 1.3 million open-access, queryable perturbation profiles and has uncovered previously unknown mechanisms of drug effects, allowing researchers to identify transcriptional responses and/or perturbagens that mimic or reverse them. Although studies have employed CMAP to identify druggable targets across multiple diseases^14^, the most commonly used version of the platform utilizes perturbation profiles generated in immortalized cell lines, limiting its ability to uncover complexities of CNS disease biology, particularly those impacted by patient genetics.

Here, we adapt the CMAP/L1000 platform to study the biology of CNS diseases such as SMA, generating a tool we term CMAP_neuro_. To this end, we differentiated moderate to severe SMA patient-specific iPSCs into NGN2 cortical glutamatergic neurons, exposed each line to a novel library of 360 neuroactive CNS disease-related perturbagens, and measured transcriptional changes in a curated set of 467 neural genes. In doing so, we generated a database of gene expression profiles depicting response of SMA neurons to diverse perturbagens, and in the process, identified compounds that reversed or exacerbated differences between severe and moderate SMA. Finally, to expand the accessibility of this resource, we generated a queryable dataset as part of CMAP’s CLUE.io portal (https://clue.io) for researchers to: 1.) identify CNS disease-relevant perturbagens that mimic or reverse a set of differentially expressed genes, or 2.) explore the transcriptional response of a given perturbation in diverse SMA neuronal cells.

## Results

### Generation and characterization of type 0, 1, and 3 SMA patient-specific iPSC-derived NGN2 neurons

To represent SMA subtypes in our CMAP_neuro_ platform, we took advantage of previously characterized iPSC lines derived from 3 type 0/1 and 3 type 3 SMA patients (**Figure 1a**)^15–18^. Included lines represent 2 female and 4 male individuals, generated from biopsies obtained from patients with varying disease severities as noted by Hammersmith Functional Motor Scale (HFSME) and Children’s Hospital of Philadelphia Infant Test of Neuromuscular Disorders (CHOP INTEND) scores (clinical metrics for SMA patients in children and infants, respectively)^19,20^.

**Figure 1.**
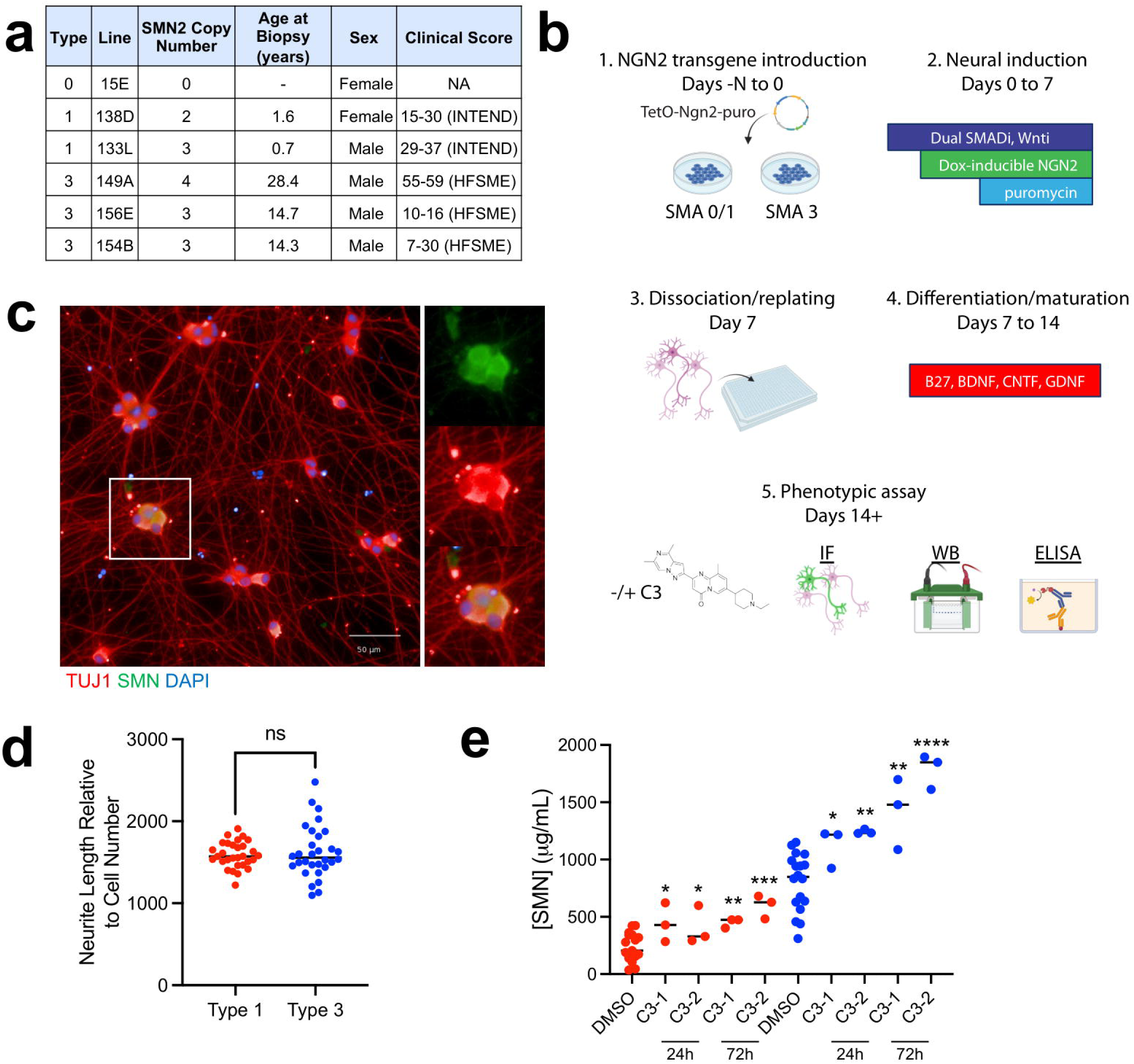
Development and characterization of SMA patient-derived NGN2 neurons. **(a)** Clinical characteristics of SMA patients from which iPSC lines were derived. **(b)** SMA patient fibroblast samples were reprogrammed into iPSCs and differentiated into NGN2 cortical-like neuronal cells via forward programming. 7-days post-dissociation and plating into 96-well plate format, cells were stained via IF or protein lysates were generated for WB and ELISA −/+ splicing modulator C3. **(c)** iPSCs successfully differentiate into cortical-like NGN2 cells that express TUJ1 and SMN by IF. Magnified inset designated by white box. **(d)** NGN2 cells differentiated from type 1 and type 3 iPSC lines exhibit equivalent neurite length as measured by TUJ1 staining. The total neurite length relative to the number of cells in each well is depicted. NS = not significant. **(e)** At baseline, type 1 iPSC-derived NGN2 cells exhibit lower SMN levels compared to type 3 iPSC-derived NGN2 cells via ELISA with detectible increases in SMN being produced by exposure to the SMN splicing modulator C3 (24h, 72h; C3-1 = 1 µM and C3-2 = 5 µM). Each condition compared to its respective DMSO control; unpaired t-test for significance; *p<0.05, **p<0.01, ***p<0.001, ****p<0.0001. Blue dots denote type 3 cells, red dots denote type 1 cells.

To generate a sufficient quantity of homogeneous, post-mitotic human neurons for perturbation-based screening, we employed a commonly used, rapid and reproducible neuronal differentiation method based on combining small molecule patterning toward a cortical excitatory identity via overexpression of the lineage-specific transcription factor Neurogenin-2 (NGN2)^21^ (**Figure 1b**). In differentiating each iPSC line into NGN2 neurons of cortical and glutamatergic identity, we observed widespread expression of SMN and pan-neuronal marker TUJ1 by immunofluorescence (IF) (**Figure 1c**). NGN2 neurons derived from moderate or severe SMA types exhibited similar cell viability as measured by normalized neurite length via TUJ1 staining (**Figure 1d**). This observation is in contrast with SMA patient iPSC-derived motor neurons, which exhibit observable phenotypic differences such as type 1 neurons degenerating more rapidly than type 3 neurons under standard culture conditions (i.e., without added stressors)^17^. However, in line with clinical observations^1^, type 1 NGN2 neurons exhibited substantially lower SMN protein levels than type 3 cells by ELISA (**Figure 1e)**. In both type 1 and 3 cells, exposure to a small molecule SMN splicing modulator C3 (a chemical analog of risdiplam) for 24 and 72 hours increased SMN protein levels by ELISA (**Figure 1e**). Aside from intracellular SMN levels, all other morphological characteristics examined were similar between all type 0/1 and type 3 cell lines tested.

### A chemical genetic screening platform to measure the transcriptional responses of SMA neurons to diverse neuroactive perturbagens

After assessing the ability of patient-specific iPSC-derived NGN2 cells to recapitulate some aspects of SMA biology, we next sought to modify and expand the CMAP platform originally focused on cancer-related immortalized cell lines to SMA NGN2 neurons (**Figure 2a**). To accomplish this, we differentiated 3 type 0/1 and 3 type 3 patient-specific iPSC lines into NGN2 neurons, matured the cells for 7 days (a total of 14 days of differentiation), and exposed each line to a curated set of 360 compounds at 2 concentrations for 24 hours. Compounds were selected based on known neuroactivity and represent diverse mechanisms of action implicated in many neurological disorders, including calcium channel blockers, glutamate receptor antagonists, acetylcholine receptor agonists, and HDAC inhibitors (**Figure 2b**, **Supplemental Table 1**).

**Figure 2.**
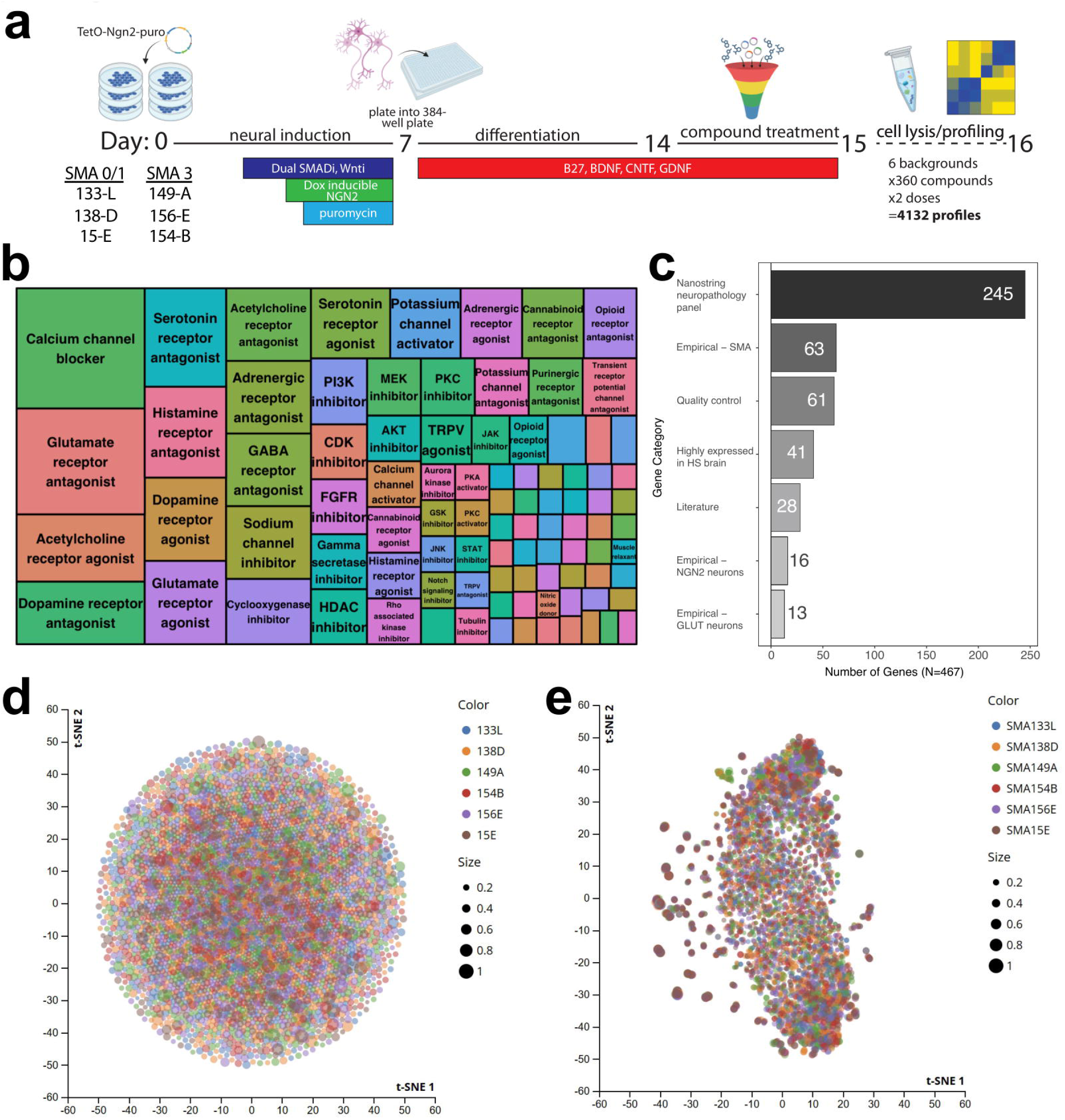
A chemical genetic platform to measure transcriptional responses to neuroactive perturbagens in iPSC-derived NGN2 neurons. **(a)** IPSCs derived from 6 SMA patients (1 type 0, 2 type 1, and 3 type 3) were differentiated into NGN2 neurons. After dissociation and plating into 384-well plates, cells were exposed to 360 compounds at 2 concentrations (0.5 µM, 5 µM) for 24 or 72 hours and subsequently lysed for targeted transcriptional profiling. **(b)** Mosaic plot depicting approximate mechanism of action for each compound included. 360 perturbagens were selected based on known activity on neurons (perturbagens noted in **Supplemental Table 1**). **(c)** Bar graph noting pathways related to genes transcriptionally profiled post-perturbation. Magnitude represents the number of genes belonging to each pathway. HS = *Homo sapiens*; GLUT = glutamatergic neurons; quality control = stress response (genes noted in **Supplemental Table 2**). **(d)** t-SNE projection of NDIS-D L1000 data show little stratification by SMA type, suggesting that type 1 and 3 SMA cells do not universally respond differently to compound perturbation. Each point corresponds to a particular compound, cell line, and dose combination, stratified by SMA cell line, with size of the dot reflective of TAS. **(e)** t-SNE projection of CMAP_neuro_ data depicting response of type 0/1 and type 3 NGN2 cells to neuroactive compounds. Each point corresponds to a particular compound, cell line, and dose combination, stratified by SMA cell line, with the size of the dot reflective of TAS.

Given the importance of cell type in mediating response to various perturbations, along with the tissue-specific nature of SMA, we sought to develop a customized Luminex gene panel tailored to measuring changes in CNS-related genes. To this end, we curated a panel of 467 neuronal-specific genes to probe within CMAP_neuro_ (**Figure 2c**, **Supplemental Table 2**). Genes were selected based on a combination of empirical analyses and literature searches using the following criteria: high expression levels in the brain or SMA-related cell types, previous implication in SMA pathogenesis, inclusion in Nanostring’s neuropathology panel, and existing L1000 landmark genes (**Supplemental Table 3**).

To detect gene-specific sequences, Luminex beads were coupled to DNA oligos complementary to each barcode used in the CMAP_neuro_ probe pool. Probe and primer design and coupling were performed as described^12^; however, because the CMAP_neuro_ probe pool contains only 467 unique genes, we were able to couple one gene’s DNA barcode to each bead color, as opposed to the 2:1 ratio used in the standard 978-gene L1000 probe pool. As a result, we circumvented the need to perform the peak deconvolution procedure used in the standard L1000 pipeline. The CMAP_neuro_ probes were validated by performing a signature analysis comparing their measured expression values with RNAseq data from a panel of 96 reference cell lines (see Materials and Methods), with 76% of probes being recalled successfully (**Supplemental Figure 1a**). Probes which appeared to fail recall were enriched for genes with generally low expression or constant expression after perturbation, suggesting that these recall failures were not due to poor probe performance (**Supplemental Figure 1b**).

### SMA NGN2 neurons exhibit changes in transcription of neurological disease-associated genes in response to CMAP_neuro_ perturbagens

In order to visualize transcriptional differences across disease types after perturbation, we first performed t-distributed Stochastic Neighbor Embedding (t-SNE)^17^ on the CMAP_neuro_ signatures resulting from exposure of each NGN2 line to the newly curated library of neuroactive compounds. Specifically, we plotted transcriptional activity score (TAS), a metric that incorporates the magnitude and consistency of a given response (see Materials and Methods). Initially, we plotted TAS post-perturbation in genes included within the original CMAP/L1000 platform. In doing so, we observed robust changes in TAS resulting from exposure to many compounds, with very little separation by disease type (SMA 0/1 vs. SMA 3; **Figure 2d**). We next sought to investigate TAS performance using our novel CMAP_neuro_ gene panel. In doing so, we observed separation of cell lines and dosing paradigms, suggesting SMA type-specific responses to the set of assayed perturbagens (**Figure 2e**). Taken together, these data highlight increased resolution in disease subtype response to perturbagens when considering a targeted panel of CNS disease-related genes.

### Querying CMAP_neuro_ to identify compounds that modulate SMA subtype-affected genes

Above, we show that by incorporating neuroactive compounds and assessing neurological disease-associated genes utilizing CMAP_neuro_, we can identify compounds that elicit strong transcriptional effects in SMA patient-specific NGN2 neurons. With those data in hand, we next sought to employ the traditional CMAP workflow to identify compounds that mimic or reverse a transcriptional signature of interest on a novel dataset – here, differences in gene expression between moderate and severe SMA subtypes.

To test the “Query” function of CMAP_neuro_, we differentiated 3 type 0/1 and 3 type 3 SMA patient-specific iPSCs into NGN2 neurons, isolated RNA from each at baseline, and performed bulk RNAseq (**Figure 3a**). In doing so, we observed clear separation of moderate and severe SMA NGN2 neurons via gene expression as noted by principal component analysis (PCA) (**Figure 3b**, **Supplemental Data File 1**) (N=383 differentially expressed genes (DEGs), p-value < 0.05). NGN2 neurons from type 0/1 and type 3 SMA backgrounds exhibited similar expression levels of pan-neuronal markers (*TUBA1A*, *MAP2*, *RBFOX3*, *STMN2*), glutamatergic neuron markers (*CAMK2A*, *SLC17A6*, *GRIA1*, *GRIN1*), and mature neuron markers (*DLG4*, *VAMP2*, *SYN1*, *SNAP25*, *SYT1*, *KCNQ2*), with limited expression of pluripotency genes (*POU5F1*, *NANOG*) (**Figure 3c**), suggesting that differences between type 0/1 and type 3 NGN2 neurons are not due to differences in differentiation capacity. Moreover, DEGs between type 0/1 and 3 neurons were related to neuronal development, cell motility, and transcriptional regulation by RNA polymerase II (**Figure 3d**, **Supplemental Figure 2**), processes found to be relevant to SMA and SMN biology^22–24^.

**Figure 3.**
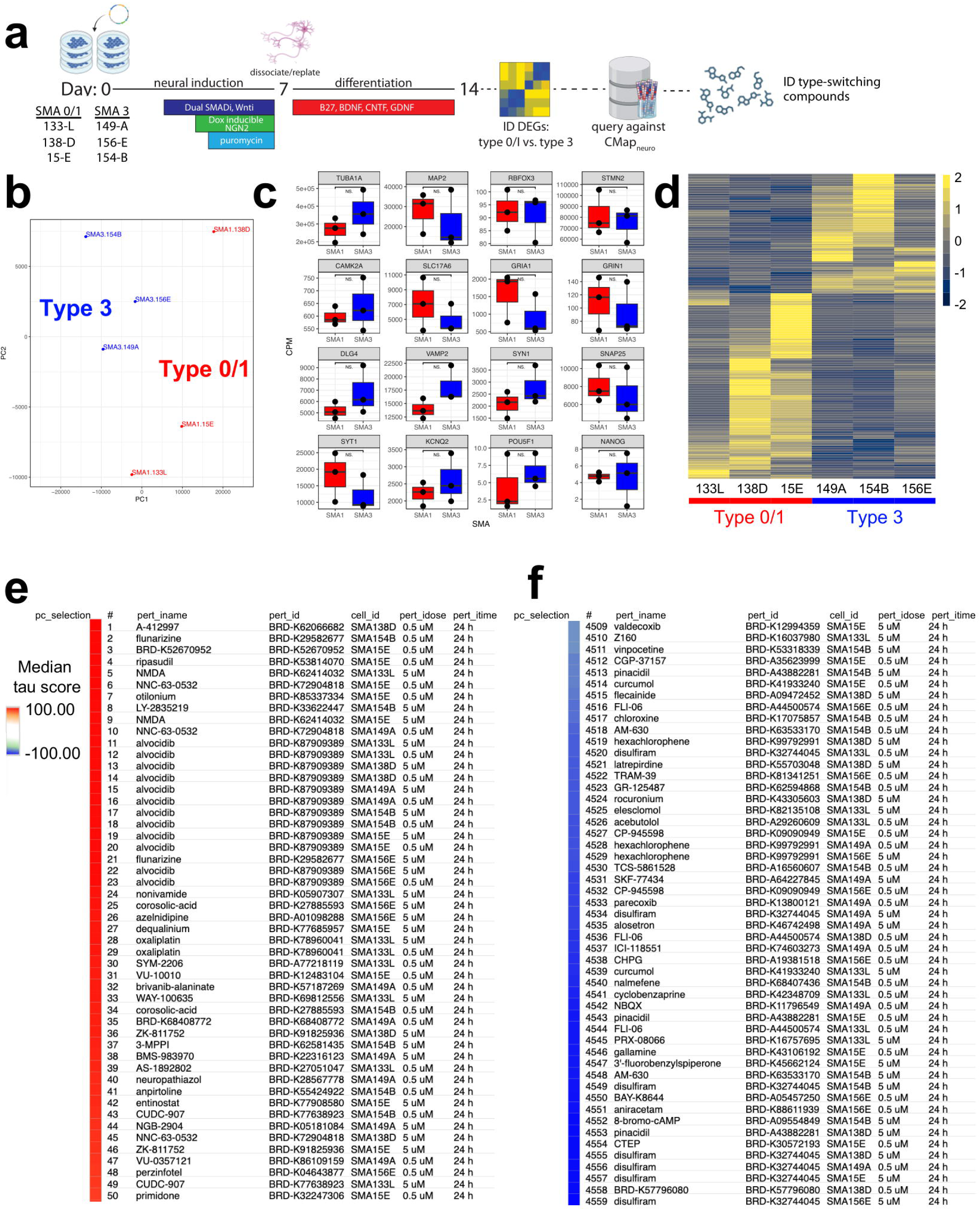
Identification of perturbagens that shift SMA type transcriptional signatures in NGN2 neurons. **(a)** Type 0/1 and type 3 SMA patient-specific iPSCs were differentiated into NGN2 neurons, RNA was isolated from each, and RNAseq was performed. DEGs were subsequently queried against the CMAP_neuro_ database to identify compounds that mimic or reverse SMA type-specific gene signatures. **(b)** Type 0/1 and type 3 NGN2 neurons form distinct clusters by PCA, separated by DEGs. **(c)** NGN2 neurons from either SMA type exhibit high expression of pan-neuronal (*TUBA1A*, *MAP2*, *RBFOX3, STMN2*), glutamatergic (*CAMK2A*, *SLC17A6*, *GRIA1*, *GRIN1*), and synaptic/functional markers (*DLG4*, *VAMP2*, *SYN1*, *SNAP25*, *SYT1*, *KCNQ2*) with minimal expression of pluripotency genes (*POU5F1*, *NANOG*). No differences in expression were observed across SMA types (Wilcoxon Rank Sum test, p < 0.05). Dots depict individual cell lines; CPM = counts per million. **(d)** Heatmap noting gene expression differences between type 0/1 (left) and type 3 (right) SMA NGN2 neurons. Each column represents NGN2 neurons differentiated from 1 SMA iPSC line. Log_2_FC depicted. **(e)** DEGs between type 0/1 and type 3 cells were input into CMAP_neuro_ and compounds that mimic (red) or reverse (blue) the signatures were identified. Median tau scores represent the strength of the connectivity across the cell lines.

We subsequently mapped the Ensembl identifiers from these data to HGNC and split the DEG list into the top 137 positive and 150 negative log2 fold-change genes to input into CMAP_neuro_. Based on median tau score, the perturbagen alvocidib positively connected with (i.e., “mimics”) gene expression differences between type 0/1 and 3 SMA cells (**Figure 3e**). Notably, alvocidib (flavopiridol) has been shown to prevent apoptosis, and has been associated with neuroprotection in cerebellar granule neurons^25^. Conversely, disulfiram negatively connected with (i.e., “reverses”) gene expression differences between types 0/1 and 3 cell lines based on median tau score (**Figure 3f**), and has been demonstrated to prevent pyroptosis^26^ and may have potential in modulating AD processes^27^. Here, CMAP_neuro_ allowed for the identification of several compounds that may impact SMA subtype-specific biology.

### An open-source queryable SMA perturbational dataset to identify compounds that mimic or reverse transcriptional states and disease associated processes

Through the CMAP_neuro_ platform, we generated a dataset of >13,000 perturbational gene expression profiles (>4,500 signatures after collapsing biological replicates) in 6 SMA lines, representing the largest perturbational transcriptional dataset in SMA cells. To maximize the utility of these data for the larger research community, we have made them available for download and interactive query analysis via https://clue.io (**Figure 4a**). Collectively, these SMA-focused datasets enable researchers to: 1.) identify compounds that mimic or reverse selected gene expression signatures, and 2.) explore the transcriptional responses of SMA neurons to given compounds.

**Figure 4.**
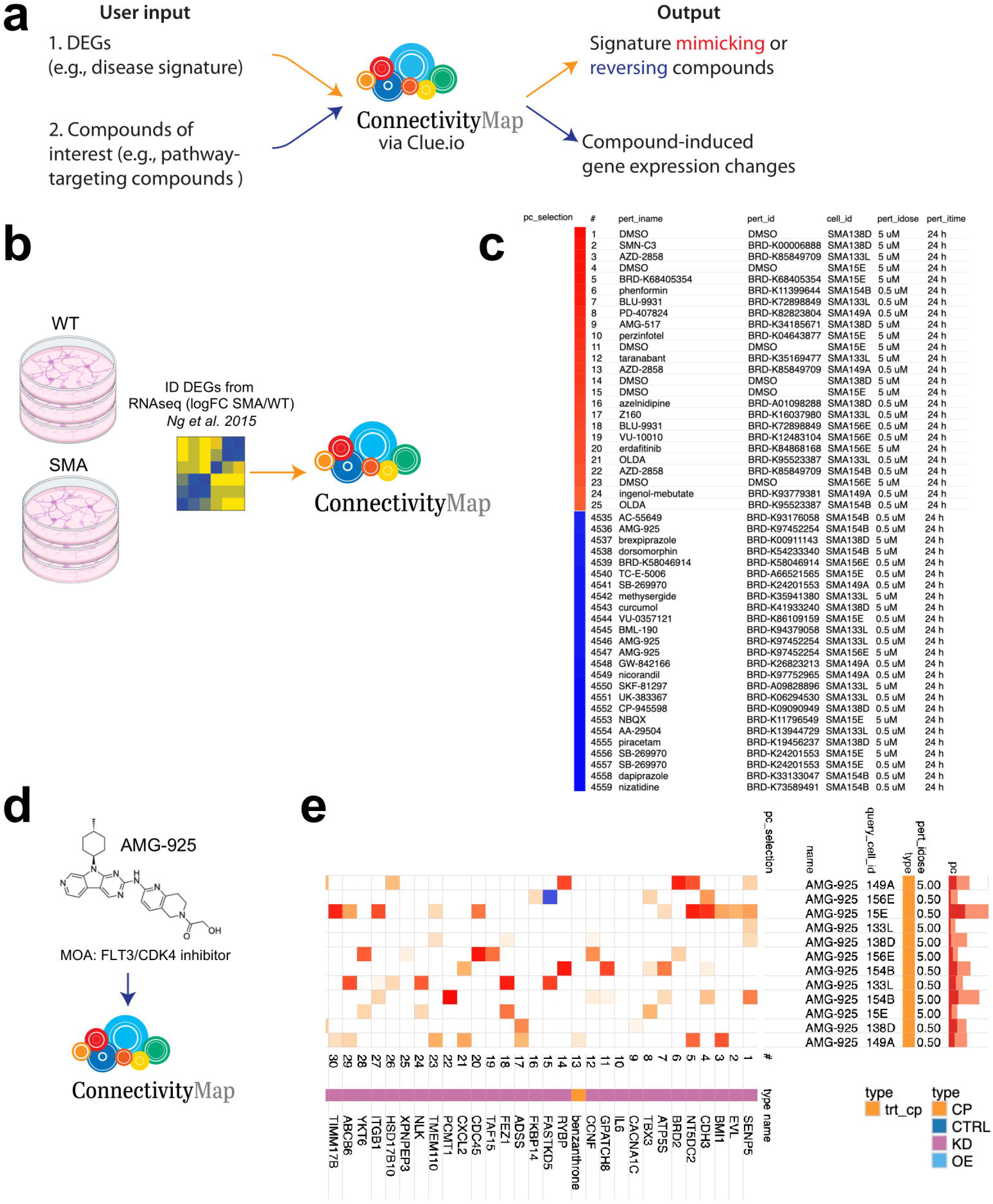
An open-source queryable platform to identify neuroactive perturbagens that induce or reverse disease relevant gene expression changes in SMA patient-specific NGN2 neurons. **(a)** Two use cases for CMAP_neuro_. In the first, users enter a list of DEGs (e.g., comparing disease to healthy cells) into the CMAP_neuro_ platform and receive a list of compounds that mimic or reverse similar DEGs in SMA NGN2 neurons. In the second, users enter a compound of interest (e.g., one associated with a given biological process) and receive genes whose expression levels are modulated in NGN2 neurons. **(b)** DEGs were obtained from Ng et al. 2015^16^ in an experiment in which wild-type and SMA iPSC-derived motor neurons were produced and subjected to RNA-seq. **(c)** Selected DEGs from **(b)** were inputted into CMAP_neuro_ and signature mimetic or reversal compounds were identified. Directionality of noted DEGs was SMA/wild-type motor neurons. Top 25 mimicking (red bar) and 25 reversing (blue bar) compounds are listed. **(d)** AMG-925, a compound found to induce opposite gene expression changes compared to inputted SMA/wild-type motor neurons from (b), was entered into CMAP_neuro_. **(e)** Top 30 perturbagens that elicit gene expression changes similar to AMG-925 in type 0/1 and type 3 NGN2 neurons are noted. pc_selection = perturbagen selected, pert_id = perturbagen name, cell_id = cell line data is received from, pert_idose = dose of compound eliciting given response, pert_itime = exposure time of compound eliciting given response, query_cell_id = cell line data is derived from in Touchstone Connectivity, pc = the percent of total perturbagens querying the row against the column, type = classification of perturbagen entered (here, AMG-925), CP = compound, CTRL = control, KD = knockdown, OE = overexpression.

To further test the applicability of CMAP_neuro_ and its corresponding L1000 datasets, we analyzed publicly available RNA-seq data comparing motor neurons differentiated from type 1 SMA and wild-type iPSCs^16^ (**Figure 4b**). To do this, we sorted a total of 913 DEGs (FDR < 0.01) by absolute log2 fold change and inputted the top 150 up and down gene symbols into both CMAP_neuro_ and the original CMAP/L1000 queries. Utilizing CMAP_neuro_, positive median tau scores (“mimic” signatures) showed overrepresentation of perturbagens such as OLDA, a combination of oleic acid and dopamine that serves as an agonist for the TRPV1 receptor that modifies locomotor behavior^28^ (**Figure 4c, top**). In other words, OLDA results in gene expression differences similar to those seen when comparing SMA to wild-type motor neurons. Conversely, the negative median tau scores (reverse signatures) showed several hits for AMG-925, a dual FLT3/CDK4 inhibitor involved in cell growth and survival being investigated for use in acute myeloid leukemia^29^ (**Figure 4c, bottom**). Here, AMG-925 results in gene expression changes that are directionally opposite to those observed between SMA and wild-type motor neurons (i.e., shifting an SMA transcriptional state toward a wild-type state).

In a second use case, the “Touchstone Connectivity” function available in the L1000 dataset allows researchers to select a perturbagen, cell line, and concentration (or multiple selections thereof) and identify genes whose expression is altered in SMA NGN2 neurons. To demonstrate this function, we selected the compound AMG-925 (found in **Figure 4c** to reverse an SMA vs. wild-type DEG signature in CMAP_neuro_). After entering AMG-925 into the Touchstone Connectivity function (**Figure 4d**), we received perturbagens that elicit similar gene expression changes to AMG-925 in type 0/1 and type 3 NGN2 neurons (**Figure 4e**). For example, in all backgrounds, genes modified via exposure to AMG-925 are similar to overall expression patterns affected by knockdown of genes involved in proliferation/regulation of cell cycle (*BRD2*, *TBX3*, *BMI1*, *ATPS5*), epigenetic modifications (*SENP5*), and cell signaling (*CDH3*, *EVL*, *CACNA1C*, *IL6*). As demonstrated here, CMAP_neuro_ represents a novel tool for researchers to identify CNS disease-related compounds and the neuro-related genes they impact.

## Discussion

Despite advances in our understanding of SMA biology, the efficacy of current therapeutic strategies relies on increasing SMN levels. Added clinical benefit may be achieved via therapeutics which improve disease-associated phenotypes even without increasing SMN levels. Due to the genetic and clinical heterogeneity of CNS disorders such as SMA, identification of such druggable targets is highly complicated. To address this, we developed the CMAP_neuro_ platform to employ chemical genetic-based screening to identify gene expression changes in SMA patient-derived neurons resulting from exposure to a compendium of neuroactive compounds. In doing so, we generated more than 4,000 transcriptional profiles representing the response of moderate and severe SMA iPSC-derived neuronal cells to diverse neuroactive compounds.

In addition to allowing insight into differences in response to perturbations across SMA neurons, CMAP_neuro_ provides a rich dataset representing transcriptional responses of cortical-like neurons to diverse, neuroactive chemical perturbagens. Similar to the existing CMAP/L1000 platform, CMAP_neuro_ represents an open-access, readily queryable dataset. Utilizing the CLUE.io application (https://clue.io), users can enter a set of genes and be directed to a list of perturbagens which elicit a similar or related transcriptional response in SMA 0, 1, or 3 neurons. Conversely, researchers can query a compound related to a particular pathway or mechanism of action to ask how its activation may affect the transcriptional profile of human SMA neurons.

Here, we have adapted the CMAP/L1000 platform, initially developed to study cancer cells, to investigate neurodegenerative disorders such as SMA via two approaches. First, the original platform utilized immortalized cancer cell lines (e.g., prostate cancer, breast cancer, cervical cancer, myelogenous leukemia, and acute promyelocytic leukemia cells)^12,13^ to generate its reference dataset. Although advantageous given the initial focus of the program, immortalized non-neuronal cell lines are often suboptimal for studying neurodegenerative disorders, neglecting the highly important genetic context of the patient and endogenous cell type. In place of immortalized cell lines, we utilized NGN2 cortical-like neurons differentiated from iPSCs derived from SMA patients. Importantly, these cells are more likely to reflect processes dysregulated in neurodegenerative disorders and incorporate the complete genome of the patient – highly important given the often unclear genotype:phenotype relationships seen in CNS disease. Second, we included a targeted library of perturbagens and a panel of genes associated with neurological tissue and disease. In accordance with our hypothesis, assessing the impact of neuro-related genes revealed greater separation across perturbagens as noted via t-SNE representation (**Figure 2e**).

In this work, we highlight the ability of the CMAP_neuro_ platform to 1.) identify compounds that mimic or reverse a given gene expression signature, and 2.) assess the impact of a given compound on the transcriptional profile of SMA NGN2 neurons. To achieve the first goal, we inputted DEGs from a previously published RNAseq analysis^16^ comparing wild-type and type 1 SMA iPSC-derived motor neurons (**Figure 4**) into CMAP_neuro_ (via https://clue.io), with the goal of identifying compounds that induce transcriptional changes seen between healthy and SMA motor neurons. Importantly, these compounds may represent a way to improve neuronal health in SMA beyond raising SMN protein levels. In doing so, we identified AMG-925, a CDK4 inhibitor, as one of the top compounds phenocopying transcriptional differences between wild-type and SMA motor neurons. While AMG-925 has not been studied in the context of SMA, its influence on cell cycle progression may impact neuronal behavior in disease, with inhibition of CDK4 previously shown to rescue toxicity in SMA neurons^30^. Building on this, we then entered AMG-925 into CMAP_neuro_ to profile gene expression changes resulting from its exposure to moderate and severe SMA NGN2 cells. In doing so, we observed changes in expression of genes associated with proliferation (e.g., *BRD2*, *TBX3*, *BMI1*, *ATPS5*) in line with its noted function. Future work may take advantage of these data by investigating the impact of AMG-925 on survival of SMA neuronal cells in various *in vitro* assays.

In a similar vein, researchers have been utilizing the original CMAP/L1000 platform to identify compounds that mimic or reverse gene expression signatures observed in a variety of disease and cell types. In one example, researchers employed CMAP/L1000 to identify compounds that reverse gene expression differences observed within muscle tissue of SMN knockout and wild-type mice^31^. In doing so, they identified several candidate compounds that induce gene expression patterns opposite of SMN^−/−^compared to wild-type mice that, upon further investigation, were found to rescue SMA-related muscle damage *in vitro* and *in vivo*. Beyond SMA, Shindyapina and colleagues^32^ recently utilized CMAP/L1000 to identify compounds that induce gene expression patterns associated with proposed biomarkers of aging and longevity. Impressively, several compounds they identified were found to increase lifespan in male mice.

The CMAP_neuro_ platform described here has the potential to identify compounds therapeutically relevant to SMA disease progression and wider CNS disorders. Despite this promise, there are several limitations to be considered regarding this assay. First, this platform is based on transcriptional profiles generated in NGN2 neurons. Although these cells are non-cancerous and more relevant to diseases of the CNS, non-neuronal cells (e.g., vasculature and glial cells) are also critical to the pathogenesis of many neurological diseases. Similarly, NGN2 neurons are closer to neuron populations found in the cortex of the brain, in contrast to spinal cord motor neurons more typically affected by SMA. Here, NGN2 neurons were selected given the robustness, relative homogeneity, and large scalability of their differentiation protocol – characteristics necessary for generating a chemical genetic-based transcriptional screening dataset. However, it is important to note that these cells did not reproduce phenotypic differences originally seen in SMA motor neurons. Improvements to CMAP_neuro_ could be made by performing bulk RNA sequencing or optical pooled screening as the output on disease-specific neurons under conditions in which perturbagen treatment produces detectable phenotypic differences. Notably, longer exposure paradigms are more likely to replicate processes dysregulated in CNS disorders that often evolve over much greater periods of time.

Despite these limitations, comprehensive use of compound-based perturbation, coupled with transcriptional screening, has great potential for uncovering novel insights into neuronal cell biology and disease processes. Importantly, though originally designed to study SMA biology, this platform can be expanded to diseases with commonly dysregulated biological mechanisms such as generalized neurodegenerative disorders.

## Methods

### Cell culture of human pluripotent stem cells

Patient-specific iPSC lines utilized were derived as previously described^15–17^. Cells were maintained in feeder-free conditions on Matrigel (VWR, Cat. No. BD35427) and cultured in STEMFLEX media (Life Technologies, Cat. No. A3349401). Work with hiPSCs was determined to be *Not Human Subjects Research* by the Harvard University Area Institutional Review Board.

### Lentivirus production and transduction of iPSCs

Lentiviral particles containing pTet-O-Ngn2-puro constructs were generated via the SuperLenti Lentiviral Packaging Mix kit (ALSTEM, Cat. No. VP100)^21^. Briefly, packaging 293T cells were seeded at 4×10^6^ cells per 100 mm plate. 2.5 ug tet-inducible NGN2 plasmid were added to lentiviral packaging mix diluted in DMEM (ThermoFisher, Cat. No. 11330057) and added dropwise to cells. Virus-containing media was collected over the next 72-hours and concentrated via addition of Lentivirus Precipitation Solution (ALSTEM, Cat. No. VC100) and centrifugations per manufacturer’s instructions.

### NGN2 cell differentiation

NGN2 cortical-like cells were generated via a two-dimensional, feeder free, cytokine-and transgene-driven differentiation protocol as previously described^21^. Briefly, iPSCs were removed via EDTA and plated at high density in STEMFLEX media supplemented with ROCK inhibitor. After 24 hours, cells were exposed to N2 basal media supplemented with SB431542 (10 µM; R&D Systems, Cat. No. 1614), XAV939 (2 µM; ReproCell, Cat. No. 04-0046), LDN-193189 (100 nM; ReproCell, Cat. No. 04-0074), and doxycycline (2 µg/mL). The following day, cells were treated with N2 basal media containing SB431542 (5 µM), XAV939 (1 µM), LDN-193189 (50 nM), doxycycline (2 µg/mL), and puromycin (5 µg/mL; Life Technologies, Cat. No. LA1113803). On day 3, cells were exposed to N2 media supplemented with B27 (1:50), doxycycline (2 µg/mL), and puromycin (1:2000). On day 4, cells were cultured in NBM basal media supplemented with doxycycline (2 µg/ml), BDNF (10 ng/ml; Miltenyi Biotec, Cat. No. 130-103-435), CNTF (10ng/ml; Miltenyi Biotec, Cat. No. 130-123-659), and GDNF (10 ng/ml; Miltenyi Biotec, Cat. No. 130-108-986). On day 5, cells were exposed to NBM basal media supplemented with doxycycline (2 µg/ml) (Sigma Aldrich, Cat. No. D9891), BDNF (10ng/ml), CNTF (10 ng/ml), GDNF (10 ng/ml), and U/FDU (10 µM).

Prior to re-plating on day 7, wells were first coated with poly-L-ornithine (1:1000) (Sigma Aldrich, Cat. No. P4957) and poly-D-lysine (1:100) (ThermoFisher, Cat. No. A-003-E) in borate buffer overnight and then coated with fibronectin (1:200) (Corning, Cat. No. 356008) and laminin (1:200) (Life Technologies, Cat. No. 23017015) for 3 hours. On day 7, cells were removed from their original plates with trypsin (0.25%) supplemented with DNaseI and plated in media containing ROCKi in 6-, 12-, or 96-well plate format. Day 7 media is composed of NBM basal media supplemented with B27, BDNF (10 ng/ml), CNTF (10 ng/ml), and GDNF (10 ng/ml). On day 8, media was completely removed and replaced with day 7 media minus ROCKi. Half-media changes were performed every 3-4 days after plating until downstream application.

N2 basal media consists of DMEM/F12 supplemented with glutamax (1:100), dextrose (0.3%), and N2 supplement (1:500). NBM basal media consists of neurobasal media supplemented with glutamax (1:100), dextrose (0.3%), MEM NEAA (1:100), and B27 (1:50).

### SMN ELISA

Protein was isolated from each well via Enzo SMN ELISA kit (Enzo, Cat. No. ADI-900-209) per manufacturer’s instructions. Briefly, lysis buffer supplemented with PMSF (1 mM), phosphatase inhibitor (Avantor, Cat. No. 78420), and protease inhibitor cocktail (Avantor, Cat. No. 87786) was added to each well and cells were incubated on ice for 30 minutes and subsequently centrifuged at 14000 x g for 20 minutes. Supernatant was stored at −80°C until used. SMN concentration was determined via ELISA per manufacturer’s protocol.

### Immunofluorescence

Cells were fixed in 4% paraformaldehyde (VWR, Cat. No. 100503-917) for 20 minutes at room temperature. Fixed cells were washed and incubated for 1 hour at room temperature in blocking/permeabilization solution (5% FBS, 2% BSA, 0.3% Triton X-100). Following blocking/permeabilization, cells were incubated with primary antibody (TUJ1 (Sigma Aldrich, Cat. No. T2200), 1:200; SMN (BD Biosciences, Cat. No. 610647), 1:150) diluted in blocking/permeabilization solution overnight at 4°C. Cells were washed and incubated with respective secondary antibody for 1 hour at room temperature. Cells were subsequently stained with Hoechst (1:2000) for 15 minutes at room temperature and imaged via the ImageXpress Micro Confocal High-Content Imaging System (Molecular Devices).

### RNA sequencing

Bulk RNA sequencing was performed using an Illumina NextSeq (v 2.1.0) sequencer, with library preparation via Illumina TruSeq RNA Library Prep Kit v2. Base calls were quality filtered and converted to FASTQ files using bcl2fastq2 (v 2.20.0.422, Illumina) using the default parameters, and resulted in an average of 25.1M paired-end reads per cell line (average of 88% of reads passing filter). Pseudo-counts were generated from the FASTQ files using kallisto (v 0.45.0, Bray et al. 2016^33^). Kallisto transcript abundance was imported and reduced to Ensembl gene identifiers via tximport (v 1.34.0). Genes were analyzed via DESeq2 (v 1.46.0), first filtering out genes with less than three counts in three samples. Apeglm shrinkage estimates were obtained (v 1.28.0). PCA and heatmap analysis (pheatmap v 1.0.12) was conducted on the DEGs (p-value ≤ 0.05). Genes were converted to HGNC via org.Hs.eg.db (v 3.20.0). The DEGs were sorted by log2 fold change, and highest positive and negative 150 genes were used as input to the Clue.io “Query” function against the SMA and L1000 datasets. Over enrichment analysis of the top 150 positive and negative log2 fold change genes was performed using MSigDB, querying the Hallmark, KEGG, Reactome, and GO Biological Processes gene sets, with an FDR cutoff of 0.05. Significant hits were visualized using ggplot2 (v3.5.1).

For the Ng et al. 2015 SMA versus control dataset, the Cuffdiff DEGs were extracted from the supplementary data. The FDR ≤ 0.01 DEGs were sorted by log2 fold change, and the highest positive and negative 150 genes were used as input to the Clue.io “Query” function against the SMA and L1000 datasets.

### Neuro probe pool validation

To assess the fidelity of the L1000-neuro probe pool, we used the probes to generate baseline L1000 gene expression profiles for 96 unique cancer cell lines. We then performed a gene recall analysis as follows:

1. We computed the gene-wise Spearman correlation between the L1000 neuro measurements and the equivalent RNAseq log2 RPKM values for the same cell lines, obtained from the cancer cell line encyclopedia (CCLE).
2. We converted the correlations to a recall rank by computing the percentage of the 22,225 genes from the CCLE dataset with a higher correlation coefficient than the matched gene.

We observed that 309 of the 409 L1000-neuro probes (76%) included in the analysis had recall ranks less than 5%, suggesting that the majority of probes’ L1000 measurements match their corresponding expression pattern in the RNAseq data. We also observed that 65% (58 of 89) L1000-neuro probes with poor recall corresponded to genes with low average expression across the 96 cancer cell lines (**Supplemental Figure 2**), suggesting that poor recall performance was generally not due to failure of the L1000 probes.

### L1000 profiling

L1000 data were generated as described in Subramanian et al.^12^. Briefly, differentiated NGN2 neurons were plated into 384 well plates, treated with compound or vehicle control for 24 hours, and then lysed. mRNA was captured from the lysates, reverse-transcribed into cDNA, and subjected to ligation-mediated amplification (LMA) with the L1000-neuro probe pool, resulting in barcoded, biotinylated PCR amplicon. The amplicon was then hybridized to Luminex beads with complementary barcodes, stained with streptavidin-phycoerythrin (SAPE), and detected on a Luminex FlexMap 3D flow cytometer, capturing bead color (i.e., transcript identity) and the fluorescent intensity of biotinylated probe.

### L1000 data processing

L1000 data were processed as described in Subramanian et al.^12^, generating 5 levels of data:

- Level 1: Raw fluorescent intensities (FI) were captured from the Luminex FlexMAP 3D scanner for each measured gene (either neuro or standard L1000 panel).
- Level 2: To account for two genes measured by each bead barcode, for the original L1000 panel, FI data were deconvoluted, extracting the median FI (MFI) for the two genes. For the neuro panel, which measures only one gene per bead barcode, the median FI value was computed across all measured beads for the given barcode to generate MFI values
- Level 3: MFI values were loess-normalized to the ten L1000 invariant gene sets within each well (same for neuro and standard L1000), and all wells on the same detection plate were then quantile normalized, resulting in each sample having the same empirical distribution
- Level 4: Gene-wise robust z-scores were then computed for each sample, with reference distribution noted as all samples on the same plate.
- Level 5: Biological replicates were collapsed using a weighted average (each replicate weighted by its average correlation to the others).

### Signature recall analysis

We extracted the level 5 signatures of 228 compounds common to both the SMA L1000 dataset and the CMap LINCS2020 release. The LINCS signatures were restricted to exemplar signatures in the core LINCS cell lines. This resulted in 1,886 signatures. For each SMA L1000 compound, we computed the Spearman correlation between its signatures and each LINCS signature using the standard L1000 gene space. We then computed the rank of the matched compound signature in the LINCS data, with lower ranks corresponding to higher correlations. We applied thresholds of TAS ≥ 0.212 to identify active SMA L1000 signatures and recall rank of ≤ 190 (in the top ∼10% of all LINCS signatures) to signify successful recall.

### Transcriptional activity score and constituent metrics

The replicate correlation, signature strength, and transcriptional activity score metrics have been previously described^12^. We have included brief summaries of each metric below.

### Replicate correlation

To assess reproducibility across replicates included in the L1000 data, we calculate the 75^th^ quantile of Spearman correlation values obtained from all possible pairwise combination of replicates across level 4 data.

### Signature strength

Signature strength is calculated via the following equations where SS=signature strength, Z=scores, and N=number of replicates.

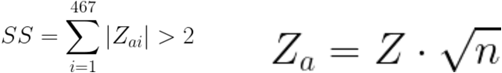

### Transcriptional activity score (TAS)

Transcriptional activity score (TAS) for a given perturbagen is calculated as follows, where SS=signature strength, CC=replicate correlation.

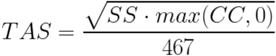

---

**Supplemental Figure 1.**
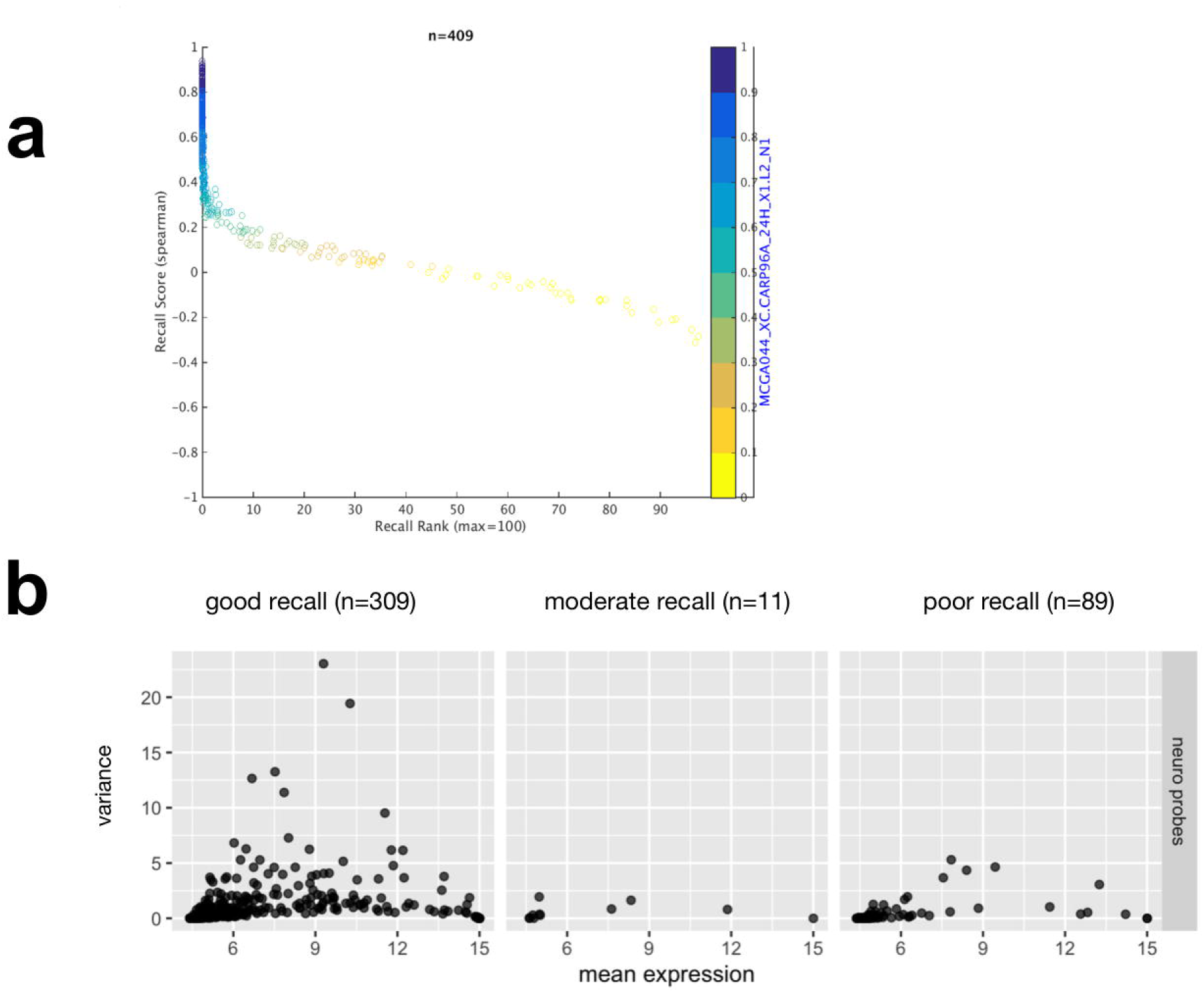
L1000_neuro_ probe pool validation. **(A)** Scatter plot depicting the Spearman correlation vs. the corresponding recall percentile rank for each of the 409 common genes. **(B)** Scatter plots of the 409 genes’ variance vs. mean expression, derived from the 96 CCLE cell lines’ log2 RPKM values, and stratified by whether the gene had good, moderate, or poor recall, corresponding to recall percentile ranks ≤ 5%, > 5% & ≤ 10%, and > 10%, respectively.

**Supplemental Figure 2.**
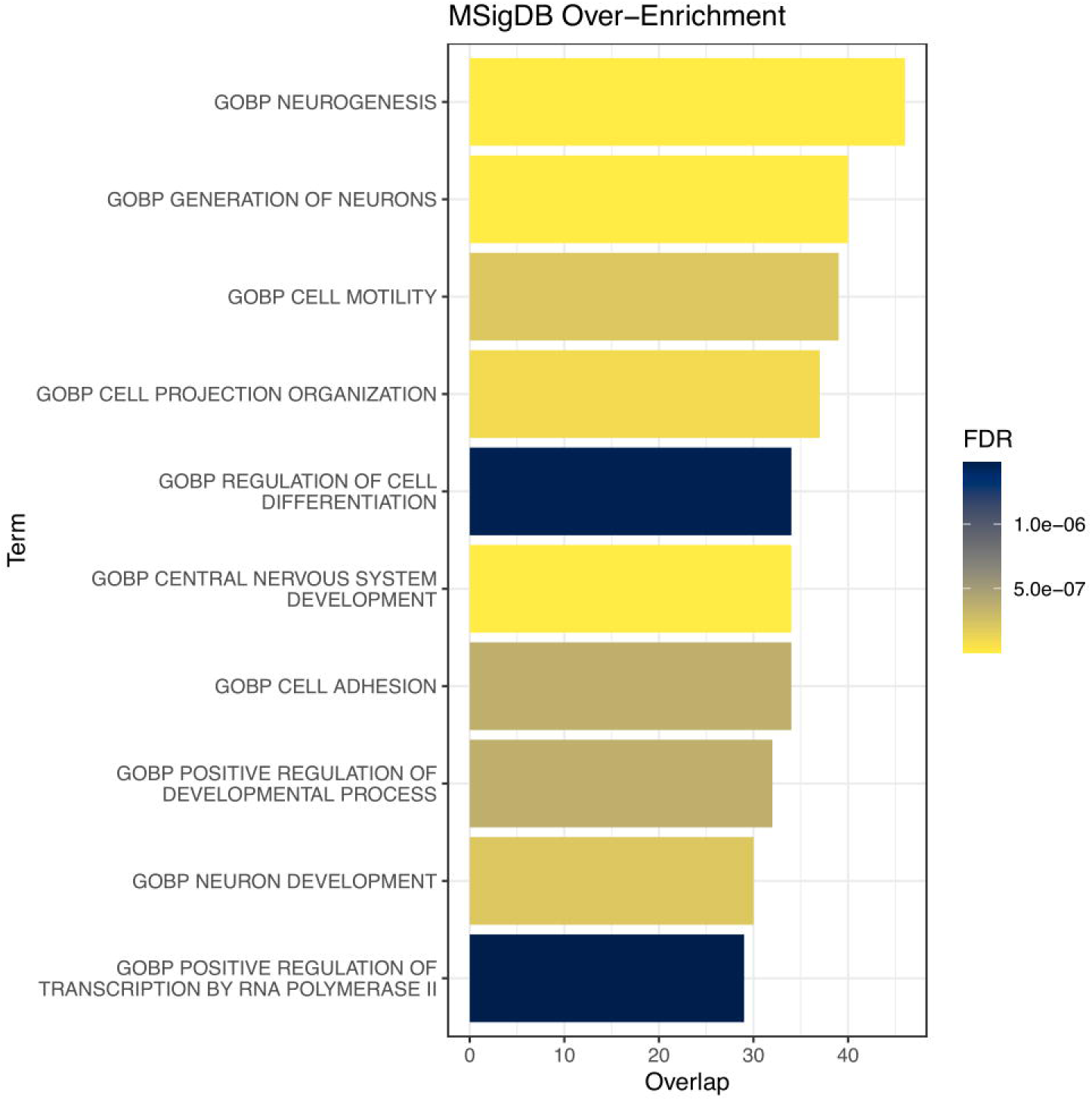
Pathway analysis of DEGs from type 0/1 and type 3 derived NGN2 neurons. MSigDB over enrichment analysis of DEGs (N=383, p-value < 0.05) from type 0/1 and type 3 SMA iPSC-derived NGN2 cells (see also **Supplemental Data File 1**). Overlap notes number of genes within each pathway noted.

**Supplemental Table 1. Perturbagens included in CMAP_neuro_ platform.**

Table lists the perturbagen id, perturbagen name, chemical formula in SMILES format, mechanism of action, and gene target.

**Supplemental Table 2. Genes assayed in CMAP_neuro_.**

Table lists the ENTREZ gene id, HGNC, and gene title for each of the L1000 n=467 CMAP_neuro_ genes.

**Supplemental Table 3. Genes assayed in L1000.**

Table lists the ENTREZ gene id, HGNC, and gene title for each of the L1000 n=978 Landmark genes.

**Supplemental Data File 1. RNA-seq analysis of iPSC NGN2 neurons.**

DESeq2 analysis of 3 SMA 0/1 and 3 SMA 3 iPSC-derived NGN2 motor neurons, with HGNC, Ensembl ID, base Mean, log 2 Fold Change, apeglm log fold change (shrinkage estimate), p-value, Benjamini-Hochberg adjusted p-value, and the normalized CPM for each sample.

## Data Availability Statement

Raw RNA-seq data will be deposited to dbGaP upon acceptance. Processed CMap data is available for interactive query and download for registered users at https://clue.io. To explore these data fully, users must create a username and password to access CMAP_neuro_ under *Neurological Disorders* within the *Data Library* tab.

## Conflict of Interest Statement

L.L.R. is a founder of Vesalius Therapeutics and VALID Tx, a member of their scientific advisory boards, and a private equity shareholder. Both are interested in formulating approaches intended to treat diseases of the nervous system and other tissues. He is also on the advisory boards of Etiome, Myrobalan Therapeutics, ProjenX and Corsalex.

## Acknowledgments

We thank Marek Orzechowski for aiding in the exploratory analysis of the CMAP_neuro_ dataset, Max Macaluso for assisting with project management and organization, and John Davis and Jacob Asiedu for assistance with maintaining the CMAP portal. Additionally, we thank Jane Lalonde and Isaac Adatto for administrative support. This work was supported by the National Institute on Aging of the NIH (1F32AG079593-01 to R.M.G., 1R01AG072086 to L.L.R.), the National Institute of Neurological Disease and Stroke of the NIH (1R01NS117407 to L.L.R.), the American Federation for Aging Research (to R.M.G.), the Simons Foundation Collaboration on Plasticity in Brain Aging to L.L.R, the Stanley Center for Psychiatric Research of the Broad Institute of Harvard and MIT, and a generous gift to the Harvard Stem Cell Institute from the Vranos Family Foundation. Co-first authors contributed equally to the manuscript, listed alphabetically here, and reserve the right to list their names first when reporting publication.

